# AlphaTims: Indexing trapped ion mobility spectrometry – time of flight data for fast and easy accession and visualization

**DOI:** 10.1101/2021.07.27.453933

**Authors:** Sander Willems, Eugenia Voytik, Patricia Skowronek, Maximilian T. Strauss, Matthias Mann

## Abstract

High resolution mass spectrometry-based proteomics generates large amounts of data, even in the standard liquid chromatography (LC) – tandem mass spectrometry configuration. Adding an ion mobility dimension vastly increases the acquired data volume, challenging both analytical processing pipelines and especially data exploration by scientists. This has necessitated data aggregation, effectively discarding much of the information present in these rich data sets. Taking trapped ion mobility spectrometry (TIMS) on a quadrupole time-of-flight platform (Q-TOF) as an example, we developed an efficient indexing scheme that represents all data points as detector arrival times on scales of minutes (LC), milliseconds (TIMS) and microseconds (TOF). In our open source AlphaTims package, data are indexed, accessed and visualized by a combination of tools of the scientific Python ecosystem. We interpret unprocessed data as a sparse 4D matrix and use just-in-time compilation to machine code with Numba, accelerating our computational procedures by several orders of magnitude while keeping to familiar indexing and slicing notations. For samples with more than six billion detector events, a modern laptop can load and index raw data in about a minute. Loading is even faster when AlphaTims has already saved indexed data in a HDF5 file, a portable scientific standard used in extremely large-scale data acquisition. Subsequently, data accession along any dimension and interactive visualization happen in milliseconds. We have found AlphaTims to be a key enabling tool to explore high dimensional LC-TIMS-QTOF data and have made it freely available as an open-source Python package with a stand-alone graphical user interface at https://github.com/MannLabs/alphatims or as part of the AlphaPept ‘ecosystem’.

**Highlights:**

- Easy visualization and fast accession of LC-TIMS-QTOF data
- Freely available graphical user interface, command-line interface and Python module on Windows, Linux and macOS.

## Introduction

The increasing amounts and complexity of data present a fundamental challenge of data accession in different scientific fields. Mass spectrometry (MS), as a leading analytical method in clinical and (bio)chemical research, is no exception. This issue is compounded when coupling MS with other techniques such as liquid chromatography (LC) and ion mobility spectrometry (IMS)^1^, which allow to efficiently separate analytes in scientific domains such as proteomics, lipidomics and metabolomics^2–4^. In our laboratory this is exemplified by time-of-flight (TOF) mass analyzers and trapped ion mobility spectrometry (TIMS)^5–7^. Typically, analytes are first separated throughout LC gradients of several minutes or hours. After ionization, they enter a TIMS tunnel where they are trapped and separated in approximately 100 milliseconds. This step discretizes continuous LC separation into ion packets with undistinguishable chromatographic retention time values and this smallest unit of LC separation is defined as a frame. After TIMS separation, a quadrupole (Q) usually provides selection for tandem MS (MS/MS) before ions reach the TOF accelerator. Ion packets are then sent orthogonally into the TOF analyzer at regular intervals of about 100 microseconds by an electrodynamic pusher. As before, such a pusher event discretizes continuous TIMS separation into ion packets with undistinguishable ion mobility (1/*K*_0_) and this smallest unit of TIMS separation is defined as a scan. Lastly, a detector at the end of the TOF accelerator discretizes continuous ion arrival times into TOF peaks of a few hundred picoseconds wide. This combination of analytical techniques, in brief LC-TIMS-QTOF, has received much attention since the introduction of the timsTOF Pro instrument (Bruker Daltonics, Germany).

The Parallel Accumulation–Serial Fragmentation (PASEF) method synchronizes ion mobility separation with quadrupole selection, combining high-throughput with high sensitivity in both data-dependent acquisition (DDA) and data-independent acquisition (DIA)^5,8^. Despite its very high data acquisition rate, the full mass resolution is maintained in MS or MS/MS mode by coupling the high-resolution TOF mass analyzer to a GHz detector. This rapid detection rate in combination with the high sensitivity often leads to billions of detector events per sample. While the actual measurements are intensity values of ion species, the exact time of a detector event can be directly converted to the TOF mass to charge (*m*/*z*), quadrupole *m*/*z*, ion mobility and chromatographic retention time values.

As a consequence of the resulting large data size, the accession and further visualization of LC-TIMS-QTOF data have proven to be challenging and slow in practice. During the last years, the single solution in the field was provided by the manufacturer’s closed-source library, integrated into Bruker’s proprietary software Compass DataAnalysis. To achieve reasonable data size and access times, this involved preprocessing steps, including data binning. However, this requires choosing parameters such as bin sizes somewhat arbitrarily and in general conceals the actual measurements. Consequently, results depend on this preprocessing and validation at the level of raw data is impractical.

Very recently, this led to parallel developments tackling some of these issues. Notable examples are OpenTIMS^9^, an open-source C++ library with bindings for the Python and R languages to read Bruker data, and MSFragger in combination with IonQuant, which allow to identify and quantify proteins rapidly without the need to preprocess raw data^10^. However, these tools were developed with specific applications in mind. We reasoned that fast and generic accession in arbitrary dimensions of the data would need to be optimized for speed, usability and extensibility. This combination would enable community-driven developments to tackle current bottlenecks such as novel implementations of feature finding algorithms, retrieval of extracted ion chromatograms (XICs) for DIA analysis or fast interactive data visualization of raw MS data.

Here we present AlphaTims, a user-friendly software tool that drastically accelerates accession and visualization of raw LC-TIMS-QTOF data compared to the vendor’s software. It provides an indexing procedure in such a way that the unprocessed data are interpreted as a sparse four-dimensional matrix. This matrix is specifically designed for LC-TIMS-QTOF data, allowing fast retrieval of arbitrary data slices along all of the available dimensions in milliseconds. It is implemented in pure Python with only a few dependencies to make it readable, flexible and lightweight. This makes it easily adoptable and adaptable by the community. At the same time, it matches the performance of programs written in the C programming language, by using the popular packages NumPy for array manipulation and Numba for just-in-time (JIT) compilation to machine code^11,12^. AlphaTims can save an indexed dataset as a single portable high-performance hierarchical data format (HDF5) file^13^, which has proven its efficiency and extensibility in various scientific fields and has also been used in MS-based proteomics before^14–16^. This further accelerates data access and allows us to store arbitrary metadata and downstream processing results. We then use Datashader, an optimized rendering Python package to plot millions of data points on standard hardware^17^, in combination with Panel and Bokeh (Python packages to build user-friendly dashboards to access and visualize data) to extend the usability of AlphaTims to a broader audience regardless of computational expertise. AlphaTims is a modular tool that is also a part of the AlphaPept^18^ (https://github.com/MannLabs/alphapept) ‘ecosystem’ developed in our department, which provides tools for the different facets of MS-based computational proteomics. It can be used as a fully stand-alone graphical user interface (GUI), command-line interface (CLI) or Python module for Windows, macOS and Linux and is freely available under an Apache license at https://github.com/MannLabs/alphatims.

## Experimental Procedures

### Sample preparation

Human cervical cancer cells (HeLa, S3, ATCC) were cultured in Dulbecco’s modified Eagle’s medium with 10% fetal bovine serum, 20mM glutamine and 1% penicillin-streptomycin (all Life Technologies Ltd., UK). Cells were collected by centrifugation, washed with phosphate-buffered saline (PBS), flash-frozen in liquid nitrogen, and stored at −80 °C.

Following the in-StageTip protocol^19^, cell lysis, reduction, and alkylation with chloroacetamide were carried out simultaneously in a lysis buffer (PreOmics, Germany). The resultant dried peptides were reconstituted in double-distilled water comprising 2 vol% acetonitrile and 0.1 vol% trifluoroacetic acid to a concentration of 200 ng/μL and further diluted with double-distilled water containing 0.1 vol% formic acid. The manufacturer’s instructions were followed to load approximately 50 ng or 200 ng peptides onto Evotips (Evosep, Denmark).

### Liquid chromatography

Purified tryptic digests were separated with either a predefined ‘200 samples per day’ (SPD) method (6 minute gradient time, 50 ng peptides) or a predefined 60 SPD method (21 minute gradient time, 200 ng peptides) on an Evosep One LC system (Evosep, Denmark)^20^. A fused silica 10 μm ID emitter (Bruker Daltonics, Germany) was placed inside a nano-electrospray source (CaptiveSpray source, Bruker Daltonics, Germany). For the 200 SPD method, the emitter was connected to a 4 cm × 150 μm reverse phase column, packed with 3 μm C_18_-beads, and for the 60 SPD method to an 8 cm × 150 μm reverse phase column, packed with 1.5 μm C_18_-beads (PepSep, Denmark). Mobile phases were water and acetonitrile, buffered with 0.1% formic acid.

Additionally, 400 ng peptides were separated over a 120 minutes gradient on a 50 cm in-house reverse-phase column with an inner diameter of 75 μm, packed with 1.9 μm C_18_-beads (Dr. Maisch Reprosil-Pur AQ, Germany) and a laser-pulled electrospray emitter. The column was heated to 60 °C in an oven compartment. The binary LC system consisted of water as buffer A and acetonitrile/water (80%/20%, v/v) as buffer B, both buffers containing 0.1% formic acid (Easy nanoLC 1200, Thermo Scientific, Germany). The gradients started with a buffer B concentration of 3%. In 95 minutes, the buffer B concentration was increased to 30%, in 5 minutes to 60%, and 5 minutes to 95%. A buffer B concentration of 95% was held for 5 min before decreasing to 5% in 5 minutes and re-equilibrating for further 5 minutes. All steps of the gradients were performed at a flow rate of 300 nL min^−1^.

### Mass spectrometry

Liquid chromatography was coupled online to a TIMS quadrupole time-of-flight instrument (timsTOF Pro, Bruker Daltonics, Germany) with ddaPASEF and diaPASEF^7,8^ via a CaptiveSpray nano-electrospray ion source. For both acquisition modes, the ion mobility dimension was calibrated with three Agilent ESI-L Tuning Mix ions (*m*/*z*, 1/*K*_0_: 622.0289 Th, 0.9848 VS cm^−2^; 922.0097 Th, 1.1895 VS cm^−2^; 1221.9906 Th, 1.3820 Vs cm^−2^). Furthermore, the collision energy was decreased linearly from 59 eV at 1/*K*_0_ = 1.6 Vs cm^−2^ to 20 eV at 1/*K*_0_ = 0.6 Vs cm^−2^.

For the ddaPASEF method, each topN acquisition cycle consisted of 4 PASEF MS/MS frames for the 200 SPD and 60 SPD methods and 10 PASEF MS/MS frames for the 120-minute gradient. The accumulation and ramp times were set to 100 milliseconds. Singly-charged precursors were excluded from fragmentation using a polygon filter in the (*m*/*z*, 1/*K*_0_) plane. Furthermore, all precursors that reached the target value of 20,000 were excluded for 0.4 min. Precursors were isolated with a quadrupole window of 2 Th for *m*/*z* < 700 and 3 Th for *m*/*z* > *700*. For diaPASEF we used the ‘high-speed’ method (*m*/*z* range: 400 to 1000 Th, 1/*K*_0_ range: 0.6 – 1.6 Vs cm^−2^, diaPASEF windows: 8 × 25 Th), as described in Meier *et al*.^8^.

A seventh sample was acquired with identical settings as the 60 SPD ddaPASEF method. To intentionally introduce anomalies, the TOF was calibrated with an offset of 1 Da and the air supply through the CaptiveSpray nano-electrospray source filter was blocked between minute 12 and 13.

### AlphaTims development

The AlphaTims source code is freely available on GitHub (https://github.com/MannLabs/alphatims) under an Apache license. The Python code (alphatims folder) is divided into two core modules: bruker.py provides the TimsTOF class and all functions to create, index and access objects from this class, whereas the utils.py module provides generic utilities for logging, compilation, parallelization and I/O. Three additional modules implement all functionality for plotting, GUI and the CLI.

In addition to the core Python code, the GitHub repository includes much introductory and background information. This includes (1) an extensive README for navigation, installation and usage instructions, (2) a Jupyter notebook folder (nbs) with a Python tutorial and a performance notebook to reproduce all timings as presented in this manuscript, (3) a documentation folder (docs) to create all documentation for the Bruker, utils and plotting modules hosted on https://alphatims.readthedocs.io, (4) a miscellaneous folder (misc) facilitating manual creation of new GUI releases and Python Package Index (PyPi) releases on https://pypi.org/project/alphatims, (5) a .github folder to perform continuous integration including testing and automatic releasing of new versions, and (6) a requirements folder to handle all dependencies.

AlphaTims is developed in pure Python and only has seven core dependencies: (1) h5py to handle HDF5 files, (2) Numba for JIT compilation, (3) pandas for tabular results, (4) pyzstd for generic decompression of Bruker binary data and (5-7) tqdm, psutil and click for CLI support. All plotting capabilities and the GUI are enabled by four additional packages: (1) Bokeh for visualizations and the dashboard, (2) hvplot to connect pandas data frames with Bokeh, (3) Datashader for fast rendering of visualizations, and (4) selenium for browser support. As an alternative to *m*/*z* and 1/*K*_0_ estimation, we also provide the option to retrieve calibrated values with Bruker libraries on Windows and Linux machines. Additional requirements files exists purely for legacy code and to facilitate development with dependencies such as e.g. pyinstaller to create the stand-alone GUI or twine to release new versions on PyPi.

### Computational system

All development and testing of AlphaTims was done on a MacBook Pro (13-inch, 2020) with a 2.3 GHz Quad-Core Intel Core i7 processor, 32 GB 3733 MHz LPDDR4X memory, and 2TB Flash storage running macOS Catalina version 10.15.7. Functionality on Linux and Windows was tested through continuous integration on default GitHub virtual machines running Ubuntu 20.04 and Windows Server 2019 (https://docs.github.com/en/actions/using-github-hosted-runners/about-github-hosted-runners).

## Results

To better explain the indexing procedure at the heart of AlphaTims, we shortly summarize the data structures used in the vendor’s software in their TIMS data format (tdf). A ‘.d folder’ contains two primary files to store raw LC-TIMS-QTOF data acquired with the timsTOF Pro (Bruker Daltonics, Germany) (**Figure 1A**). The first of these is the analysis.tdf file, an ordinary SQLite database that contains all metadata from the acquisition. It furthermore stores summarized information for each individual frame (ion packet with the same retention time values) and, if applicable, at which scans (ion packet with the same ion mobility values) the quadrupole isolation window was changed. The second file, analysis.tdf_bin, contains all raw detector events and their intensity values as compressed binary data.

**Figure 1.**
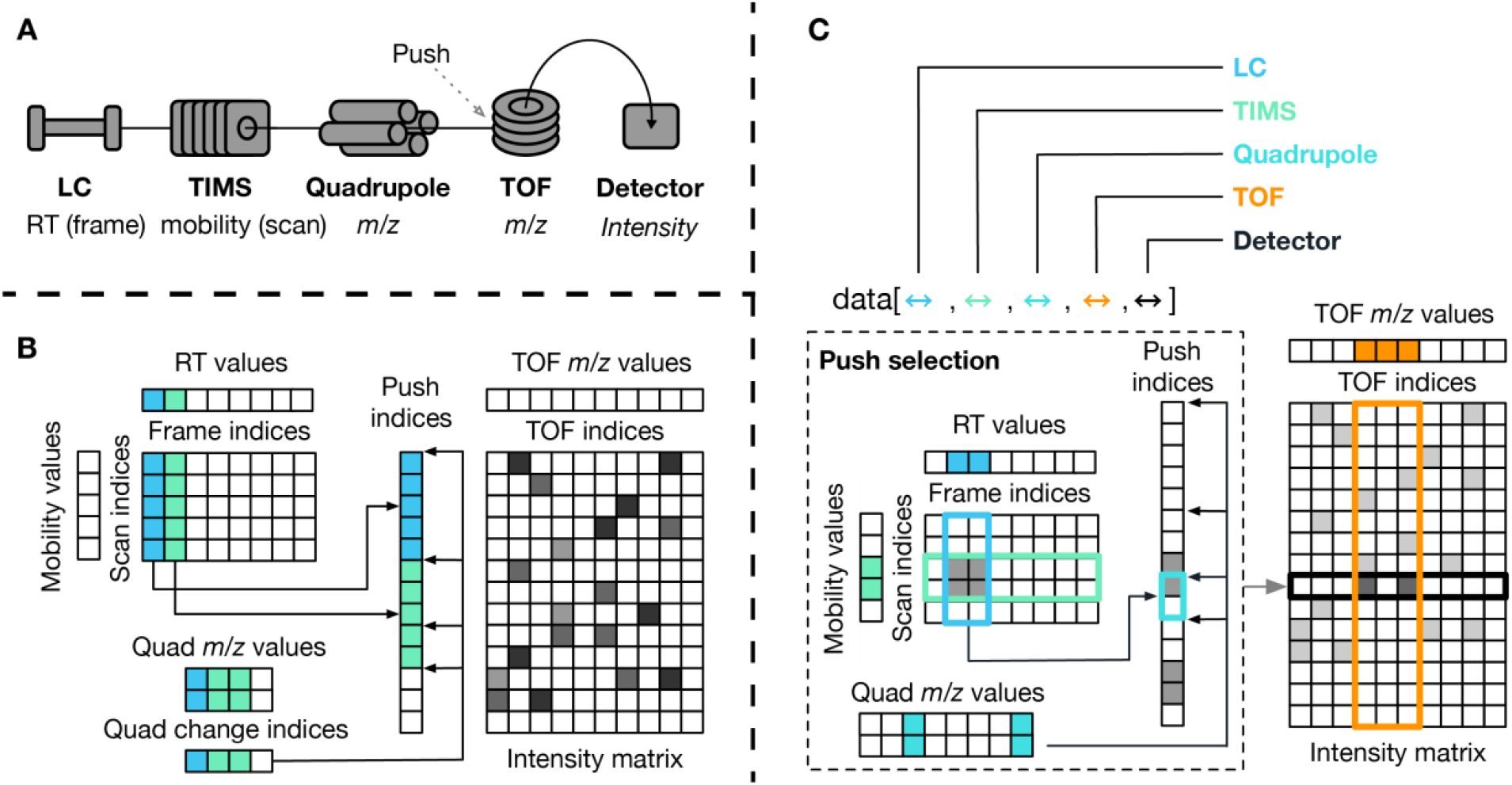
Schematic of AlphaTims’ indexing and data accession. (**A) Data dimensions:** The timsTOF instrument acquires detector events after separation and selection in four different dimensions. After passing through the LC, TIMS and quadrupole, an ion beam enters the TOF accelerator where a pusher event (synchronized with the LC, TIMS and quadrupole) sends ions in an orthogonal direction towards the detector. Liquid chromatography (LC), trapped ion mobility spectrometry (TIMS) and time-of-flight (TOF) coordinates can be represented as discrete indices (frame, scan and TOF indices) or as continuous values (RT, ion mobility and TOF m/z values). **(B) Indexing procedure:** AlphaTims uses several arrays to store LC-TIMS-QTOF data. First, the intensity values are stored in a compressed sparse row matrix (intensity matrix) with TOF indices as columns and indices of pusher events as rows (push indices). Each unique pusher event corresponds to a unique combination of a frame and scan index, according to the formula push_i_ = scan_n_ + frame_m_ · #scans. Note that the scan-frame matrix presented here is purely a visual aid and is not stored explicitly, as the unique relationship between frame, scan and push indices makes this redundant. An additional sparse array stores the push indices where the quadrupole settings are changed (quad change indices). For instance, in the first frame (blue) the quadrupole is not changed, whereas it is changed once the second frame (green) starts and another time within this frame (e.g. diaPASEF with two windows per frame). An array of equal length denotes which m/z values (lower and upper bounds) are selected with the quadrupole at each of these indices. **(C) Accession procedure:** Data accession with AlphaTims can be performed in any dimension. This can be done by providing ranges of interest either as indices or as values. In case of the latter, LC, TIMS and TOF values are always converted to the closest index by fast binary searches in their corresponding arrays. All of the selected LC and TIMS indices are then converted to push indices by the formula push_i_ = scan_n_ + frame_m_ · #scans. Since the quadrupole m/z array is not ordered, a linear pass over all quadrupole m/z values is required to determine which quadrupole index pointers are valid and only those that overlap with the previously selected push indices are retained. For each individually selected push index, a binary search retrieves all TOF indices that satisfy the requested TOF range. Finally, all selected detector events are filtered with a single pass over their corresponding intensity values to obtain the final set of detector events that satisfies the multidimensional range of interest.

### Indexing procedure and performance

AlphaTims represents relevant data from a ‘.d folder’ in multiple NumPy arrays. First, it decompresses the binary analysis.tdf_bin file to read all detector events and corresponding intensity values. While Bruker stores detector events and intensity values in a single homogeneous array, AlphaTims separates them into three distinct arrays. In the first, the (non-zero) intensity values of all detector events are stored in order of their acquisition time. A second array of equal length then stores their TOF indices as offsets for each individual pusher event. To indicate when pusher events happened, AlphaTims defines a third dense array which stores the number of detector events that are registered per pusher event. By taking the cumulative sum of this latter array, pointers are created to indicate the start and end indices of individual pusher events in the two former arrays. Together these three arrays unambiguously define a compressed sparse row matrix^21^ with indices of pusher events as rows, TOF indices as columns and intensity values as values (**Figure 1B**).

Next, AlphaTims retrieves the unique number of frame, scan and TOF indices from the analysis.tdf SQL database and estimates their respective retention time, ion mobility and TOF *m*/*z* values based on the start values, end values and array length. On Windows and Linux, this estimation of ion mobility and *m*/*z* values can also be replaced by using calibrated arrays from Bruker libraries that are integrated into AlphaTims. As there are typically 600 frames per minute, 1000 scans per frame and 400,000 detector events per pusher event, the size of these three arrays are neglectable in size compared to the total number of detector events which frequently surpasses a billion.

Lastly, another sparse array is created to indicate at which push indices the quadrupole settings change. In ddaPASEF this happens on average ten times per frame to select different precursors. In diaPASEF this depends on the acquisition scheme and desired cycle time. Typically, each frame of a recurring diaPASEF acquisition cycle is split up into eight window groups that all have different quadrupole settings. This array of quadrupole change indices is accompanied by two other arrays of equal length. The first of these is two-dimensional and defines the lower and upper quadrupole *m*/*z* values selected by the quadrupole. The second defines the precursor index. For DIA, the precursor indices are equal to the diaPASEF window group.

AlphaTims collects all these arrays, together with global and frame-specific metadata from the analysis.tdf file, and stores this as an alphatims.bruker.TimsTOF object into working memory. Since a single detector event takes up 6 bytes (an uint32 for the TOF index and an uint16 for the intensity) and their respective arrays generally dwarf all others, the required working memory (in gigabytes) is roughly equal to six times the number of detector events (in billions). The alphatims.bruker.TimsTOF object acts as a fully indexed sparse four-dimensional matrix with associated metadata. To facilitate fast reuse of this object and avoid recreation of the indices, it can be stored on disk as a portable HDF5 file with Python’s h5py package. By default, the HDF5 file size is equal to the required working memory, but compression can be used to decrease this roughly two-fold. While compression slows down loading and saving of HDF5 files approximately two to ten times, an AlphaTims object in working memory is always decompressed and interactive accession is thus unaffected. Also note that a compressed HDF5 file can always be (de)compressed and resaved, making it ideal for file transfer or archiving.

To assess the performance of AlphaTims’ indexing procedure, we acquired HeLa samples with gradients of 6, 21 and 120 minutes in both ddaPASEF and diaPASEF modes (**Experimental Procedures**). At the shortest time dimension, a single pusher event could record almost 400,000 TOF detection events in an *m*/*z* range of 100 – 1700 Th. Separation in the TIMS tunnel lasted 100 milliseconds and is composed of 1000 of these pusher events, covering a 1/*K*_0_ range of 0.6 – 1.6 Vs cm^−2^. Up to 240 billion events could thus have been recorded per minute, however, in practice no run acquired more than 0.03% of these potential detector events and the data can be considered sparse (**Figure 2**).

**Figure 2.**
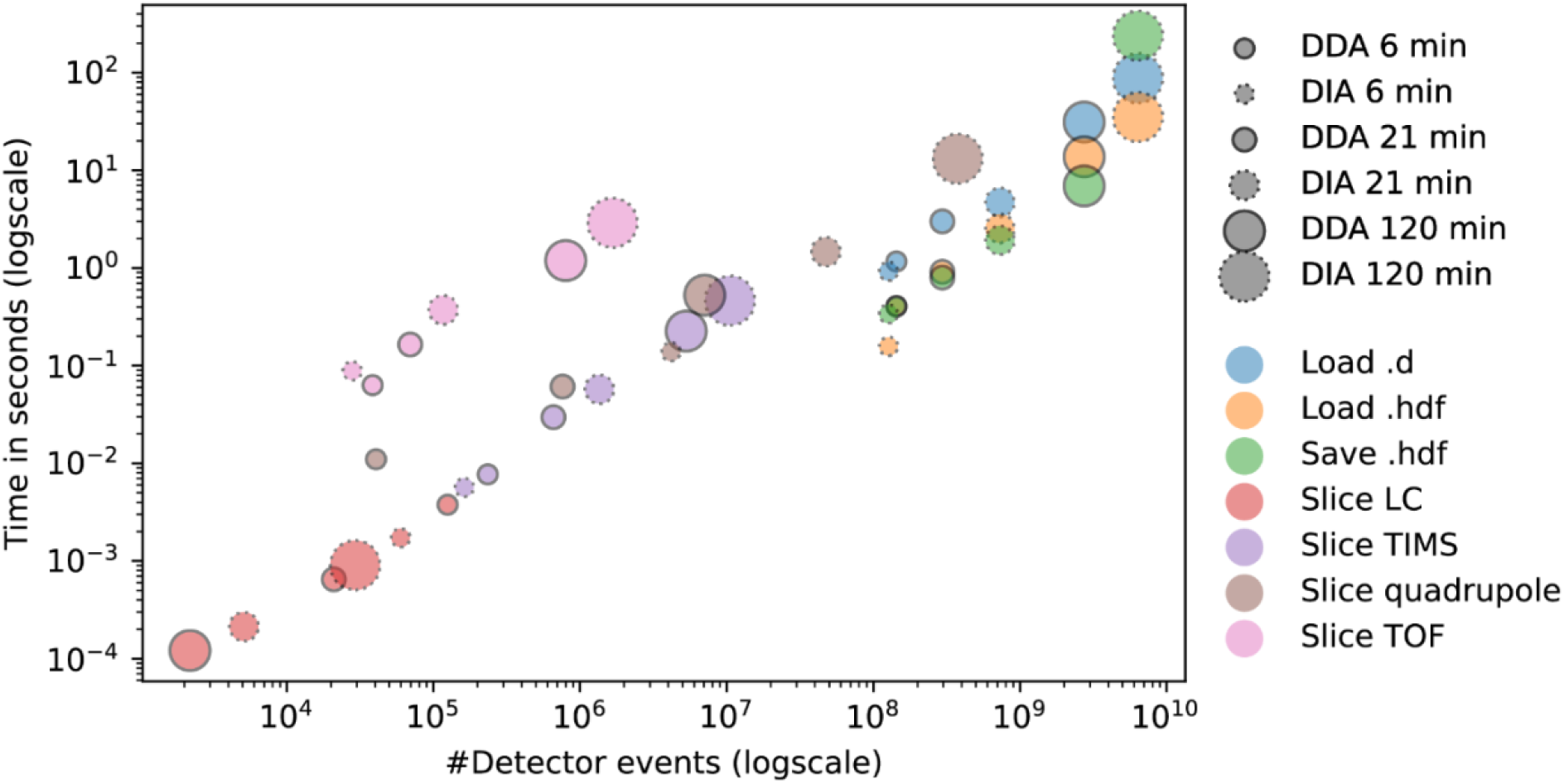
Time performance of AlphaTims. Different HeLa samples were acquired in both ddaPASEF (**full outline**) and diaPASEF (**dotted outline**) with gradient lengths of 6, 21 and 120 minutes (**Experimental Procedures**). When a raw Bruker ‘.d folder’ is read, AlphaTims needs to decompress, import and index all detector events (**blue**). Once this is done, the indexed dataset can be saved as an HDF5 file **(green)**. When an HDF5 file is read instead of a raw Bruker ‘.d folder’, no decompression or indexing is required **(orange)**. Multiple detector events of each run were retrieving by slicing each dimension individually. The retrieved detector events correspond to an LC slice with 100 ≤ retention time (s) < 100.5 **(red)**, a TIMS slice with scan index = 450 **(purple)**, a quadrupole slice with 700.0 ≤ quad m/z value < 710.0 **(brown)**, and a TOF slice with 621.9 ≤ TOF m/z value < 622.1 **(pink)**. All timings were obtained with Python timeit function for robust and reproducible results that were averaged over at least seven repeats. See https://github.com/MannLabs/alphatims/blob/master/nbs/performance.ipynb for exact numbers.

On a laptop (**Experimental Procedures**), reading all detector events into working memory and indexing them took AlphaTims less than a second for the smallest run and less than ninety seconds even for the largest run with 6.4 billion detector events. In contrast, opening any of these runs with Bruker’s Compass DataAnalysis software (v5.3) required at least double the time on a Windows desktop with overall better specifications. To speed up data import even further and allow modification or addition of downstream results, AlphaTims also allows to export the indexed data as a portable HDF5 file, which only takes seconds. When these HDF5 files are imported, no decompression and indexing is required, making them roughly three times faster to load than raw Bruker ‘.d folders’. For each of these loading and saving steps, the required time is approximately linear in function of the number of detector events and independent of LC gradient or acquisition scheme.

### Accession procedure and performance

Once data is imported and indexed, an alphatims.bruker.TimsTOF object can be accessed in all dimensions with traditional Python slices or ‘fancy index slicing’ from NumPy^12^ (**Figure 1C**). The order of the dimensions in such an object is equal to the order of their respective components in the timsTOF Pro: LC, TIMS, quadrupole, TOF and detector. Typically, the user defines a range of interest, which is translated into a slice with a single index or by a (start, stop) tuple. When decimal values are provided for the LC, TIMS or TOF dimension instead of indices, AlphaTims always assumes them to represent retention time, ion mobility or TOF *m*/*z* values. By default, these are converted to the closest integers representing frame, scan, or TOF indices by looking them up in their appropriate arrays with a fast binary search. In the case of quadrupole *m*/*z* values, precursor indices or intensities, no translation is necessary.

Once a multi-dimensional slice of interest is defined, AlphaTims first selects all the possible push indices that satisfy the LC and TIMS dimension and converts these to push indices with the formula *push_i_* = *scan_n_* + *frame_m_* · *#scans*. As these push indices are ordered, they are located in the quadrupole change index array in a single iteration. Only those push indices with a valid quadrupole *m*/*z* value are selected and for each of them appropriate TOF indices are retrieved from the sparse intensity matrix. As the TOF indices are ordered per individual pusher event, a binary search quickly retrieves all TOF indices that satisfy the requested TOF slice. Lastly, it is checked which of all the selected detector events have an intensity value that satisfies the detector slice. The results are then returned as a pandas^22^ data frame whose columns describe all indices and values, or -if desired-as a NumPy array with indices of detector events.

For each of the six HeLa samples (**Experimental Procedures**), we tested four different slices: an LC slice with retention time values between 100 and 100.5 s, a TIMS slice with a scan index of 450 providing all mass spectra at the corresponding ion mobility, a quadrupole slice with only fragments from a precursor range between 700 and 710 Th, and finally a TOF slice with *m*/*z* values between 621.9 and 622.1 (**Figure *2***). As expected, samples with longer gradients, and thus more detector events, also yield more detector events when sliced in the TIMS and TOF dimensions. While this is also true for the quadrupole dimension, the effect of being a ddaPASEF or diaPASEF method is stronger than the gradient length in these examples. This is not surprising, since the quadrupole selected just 2 or 3 Th in ddaPASEF, whereas the selected windows in diaPASEF were always 25 Th.

Next, we evaluated the time that was needed to access all of the previous data slices with AlphaTims. Due to the indexing structure, the index of any pusher event can be converted to a frame and scan index with a simple linear formula and vice versa (**Figure 1C**). As such, it can be expected that accession in these dimensions should be very fast as no actual searching is involved. Indeed, even retrieving five million detector events with slicing in the LC or TIMS dimension is done in just 0.2 seconds (**Figure *2***). Moreover, the time required to slice in these dimensions only depends on the number of detector events that are retrieved and only indirectly on the gradient length or acquisition scheme. Slicing in the quadrupole dimension is very similar. While slightly slower than the LC or TIMS dimension, there is a comparable linear dependency for the required slicing time that is purely a function of the number of detector events that are retrieved. This slowdown is due to additional filtering of quadrupole change indices from the sparse array. As this quadrupole index pointer array itself is very sparse (on average 1% non-zero elements when compared to the number of pusher events), the impact of this additional filtering is small. However, slicing in the TOF dimension is roughly an order of magnitude slower than slicing in any other dimension, primarily caused by the fact that every pusher event needs to be filtered individually, as the TOF dimension is indexed per pusher event. When TOF slicing is combined with other dimensions, fewer selected pusher events are selected which makes even this slowest step instantaneous to the user. As the time required for TOF slicing is still linearly dependent only on the number of retrieved detector events, AlphaTims is very scalable even to long gradients, very complex samples and data acquisition schemes.

### Using AlphaTims

AlphaTims is freely available as an open-source Python package with an Apache license on Windows, macOS and Linux. To enable usage for a wide audience regardless of computational background, it can be operated in any of three following modes: a stand-alone GUI, a stand-alone CLI or directly as a Python module.

#### GUI mode

A simple installer for the AlphaTims GUI can be downloaded from our GitHub page, requiring just a few mouse clicks. Both the installation and usage of AlphaTims have been made as intuitive as possible, but a comprehensive GUI manual is also available with in-depth step-by-step explanations and screenshots.

The GUI allows interactive exploration of unprocessed LC-TIMS-QTOF data conveniently in the browser. It was programmed in pure Python and uses only a few libraries of Python’s Holoviz visualization ecosystem. These include Holoviews itself and Bokeh to visualize different plots such as the total ion current (TIC), Datashader for fast rendering of these plots and Panel to combine the plots with control widgets into an interactive dashboard (**Experimental Procedures**). With the control widgets the user can slice the data simultaneously in multiple dimensions as described before (**Accession procedure and performance**). The selected coordinates can then be projected on either a single axis to show mass spectra, ion mobilograms or XICs or on multiple axes to create heatmaps in the LC, TIMS and TOF dimension.

Having reduced the visualization of LC-TIMS-QTOF to a fast and straightforward task, it can be incorporated in a wide variety of practical applications. In the following, we demonstrate this on the example of visual quality control. For this purpose, we intentionally acquired a sample with a few anomalies (including a large offset of the mass scale and temporary pressure change in the CaptiveSpray source) to see if we could indeed quickly detect any issues. There were 0.7 billion detector events in this 21-minute ddaPASEF run. The data could be imported with a single mouse click and the TIC was visible within ten seconds of opening the AlphaTims GUI. This immediately revealed an anomaly, namely the drop in ion current between minute 12 and 13 that we had engineered beforehand (**Figure 3**). Without having done any processing at all, the user is forewarned about unreliable intensity values in that region. As an important quality metric, the user can then assess the stability of added calibrant ions (1222.0 Th, 1.38 Vs cm^−2^), which is expected to be continuously present throughout the whole run. By modifying just two values of the TOF widget, we selected all ions in the *m*/*z* region between 1221.0 and 1225.0 Th. By adjusting the heatmap axes to show chromatographic retention time values on the *x*-axis and *m*/*z* values on the *y*-axis, we expect to see a continuous signal throughout the whole gradient for the calibrant spray with an *m*/*z* value of 1222.0 Th. However, there is a continuous and steady signal for an *m*/*z* value of 1223.5 Th instead, accompanied by a less intense isotope at 1224.5 Th. Based on these observations, we deduce that the TOF *m*/*z* values are greatly miss-calibrated (as intended for this sample) and that the reported *m*/*z* values are too unreliable for further analysis. Next, we changed the *y*-axis of the heatmap to show ion mobility values and inspect the detected ion at 1223.5 ±0.1 Th during the complete LC gradient. This clearly revealed another issue between minute 12 and 13. Normally the ion mobility value of the calibrant spray should remain constant at a value of 1.38 Vs cm^−2^, but in this case the apparent value drops to 1.1 Vs cm^−2^ for a full minute (as a result of the purposely altered gas flow). This coincides with the previously detected drop in the TIC, meaning that not only the intensity but also the other coordinates are unreliable in this timeframe. Thus, a brief assessment of the data in less than thirty seconds with just a few user inputs already detected and pinpointed the main issues with data quality. Other quality assessments to analyze e.g. fragmentation efficiency of ddaPASEF samples or positioning of quadrupole selections in diaPASEF samples do not require much more effort and quickly become routine even for inexperienced users.

**Figure 3.**
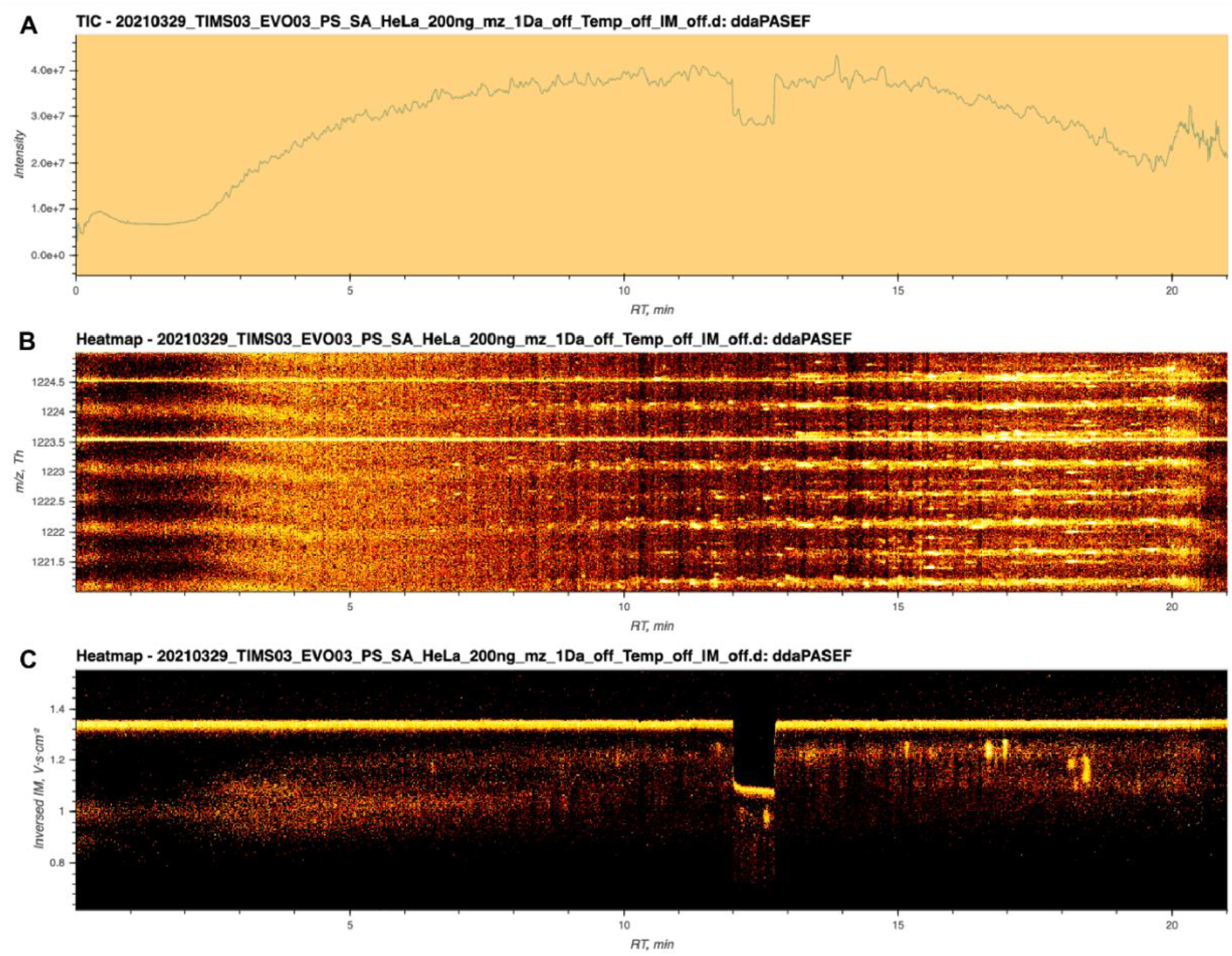
Quality control with the AlphaTims graphical user interface. **(A) Total ion current:** After importing a sample, the total ion current (TIC) is immediately available without requiring any additional user input. In this case, a clear drop in intensity between minute 12 and 13 is visible. **(B) Time of flight calibration:** By adjusting the time-of-flight (TOF) selection and plot axes widgets, the expected m/z value of a calibrant spray is visualized throughout the whole gradient. The expected value of 1222.0 Th is not present, but instead a value of 1223.5 Th is displayed. **(C) Ion mobility spectrometry stability:** When the TOF selection is narrowed to 1223.5 ± 0.1 Th and the y-axis is changed to 1/K_0_ values, a discontinuity in ion mobility is detected between minute 12 and 13.

#### CLI mode

While very easy to use, AlphaTims’ GUI requires manual input for visualization. For users who wish to automate repetitive tasks, the AlphaTims CLI provides the same functionality as the GUI. Instead of manually updating control widgets, all settings and values can be provided to the command-line either directly or with a simple script. As there is no need to display an interactive dashboard, this mode is even faster and more versatile than the GUI. More complex data slices can be selected than with the GUI, while all results can still be exported. This includes visualizations in png, or html format, csv tables with selected ion coordinates and alternative formats of the whole sample such as portable HDF5 files and mascot generic format (MGF) files. All of these commands and their options are fully documented in the CLI and a brief tutorial is available on GitHub.

#### Python mode

Even though the CLI is more flexible than the GUI, it is impossible for us to implement all imaginable use cases of AlphaTims. Instead, we also make it available as a Python module and leave it to the end user to implement any additional functionality or incorporate it into other Python projects. AlphaTims can be installed from PyPi as a Python module with the standard pip module of Python 3.8. There is both a lightweight version available with just a few dependencies that purely focuses on data indexing and accession, as well as an extended version with more dependencies that includes the complete visualization library as used for the GUI and CLI.

Enabling AlphaTims in other Python scripts or Jupyter notebooks requires a single line of code that imports the module. Some convenience functions enable logging or set the number of available threads for multithreading and ensure transparent, reproducible and efficient usage of AlphaTims. All functions of AlphaTims are implemented in pure Python and fully documented to facilitate flexibility, readability and usability. However, functions that are computationally intensive have been decorated with Numba to use just-in-time (JIT) compilation to machine code. This enables performance similar to the fastest low level languages such as C.

Importing and indexing data is done with a single command that returns an alphatims.bruker.TimsTOF object, which can be treated as a four-dimensional matrix. Inspired by the slicing approach in NumPy, one of the fundamental Python libraries for scientific computing, AlphaTims provides slicing in multiple dimensions simultaneously as described before (**Accession procedure and performance**). As a result, AlphaTims data slices can take advantage of the vast amount of Python packages that act on pandas data frames as well.

To demonstrate basic usage of AlphaTims in Python, we have provided a brief Jupyter notebook tutorial on GitHub (https://github.com/MannLabs/alphatims/blob/master/nbs/tutorial.ipynb). This notebook explains how to: set up AlphaTims and enable logging for transparent and reproducible data analysis, import samples and export indexed HDF5 files for faster reanalysis, select individual data points and data slices, and visualize data to create similar plots as with the GUI or CLI. The final part of the tutorial includes an example to show how AlphaTims can be used to investigate a specific peptide in diaPASEF data based on a spectral library created with for instance AlphaPept, Skyline or Spectronaut^18,23,24^ (**Figure *4***).

**Figure 4.**
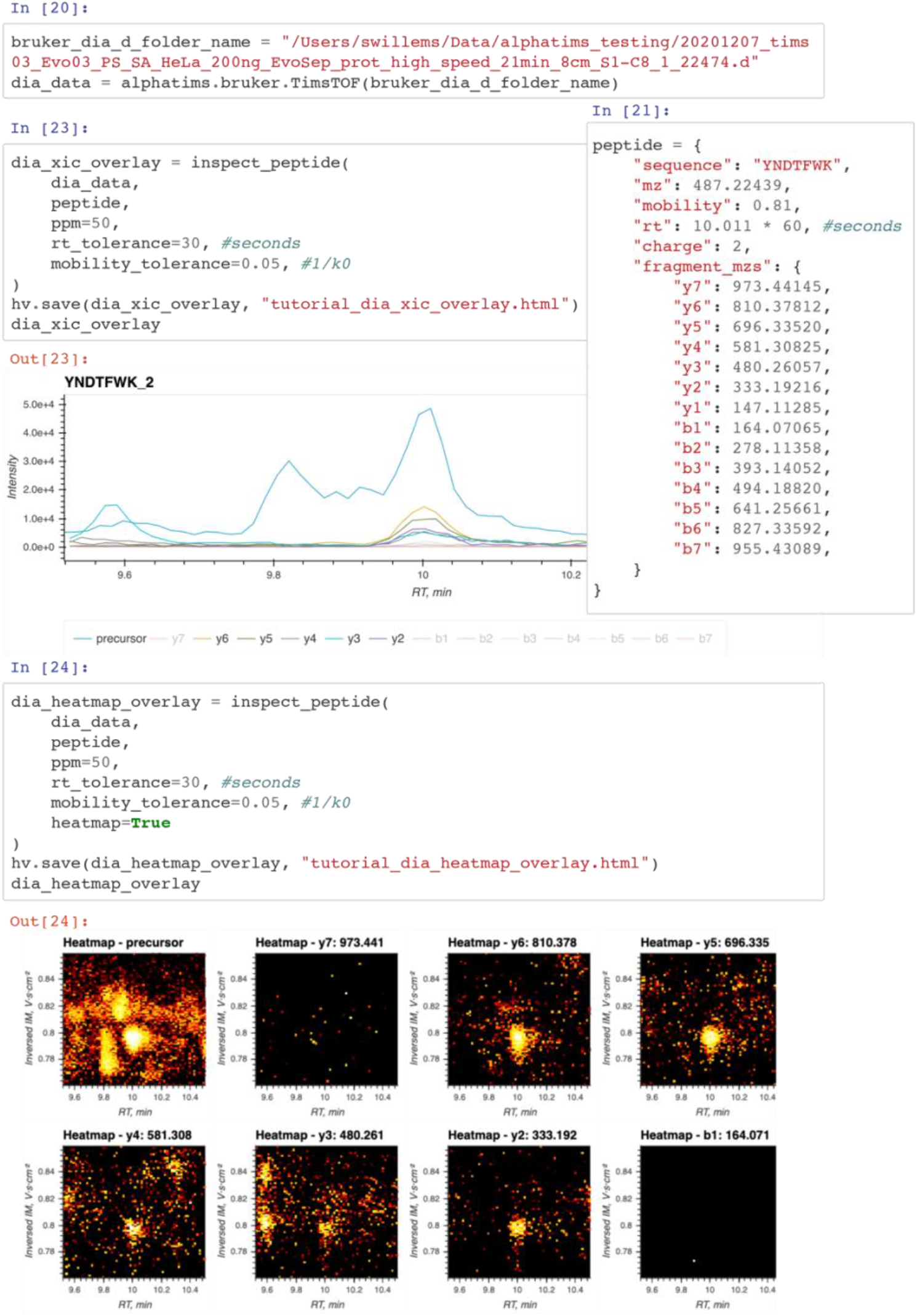
A section of a Jupyter notebook using AlphaTims as a Python module. Importing data with AlphaTims’ Python module is done with just a single command **(cell 20)**. Here, a diaPASEF sample was imported. The same sample was also acquired in ddaPASEF and a spectral library was generated with AlphaPept. Relevant coordinates of the peptide YNDTFWK were retrieved from this spectral library and passed to Python **(cell 21)**. A function ‘inspect_peptide’ was defined in Python (cell 22, see AlphaTims’ Python tutorial at https://github.com/MannLabs/alphatims/blob/master/nbs/tutorial.ipynb), allowing to visualize extracted ion chromatograms (XICs) for the doubly charged precursor and all fragments of this peptide **(cell 23)**. Based on the these XICs, some interference seems to be present for the precursor signal of this peptide. However, when the precursor and fragments of this peptide are visualized as a heatmap in both the LC and TIMS dimension, it becomes clear that this interference is fully resolved in the TIMS dimension **(cell 24)**.

The above example illustrates a use case of AlphaTims in Jupyter notebooks that have become a standard in modern data science. AlphaTims and Bruker diaPASEF data are first imported, and then all coordinates of both the precursor and all fragments of a specific peptide are defined. With a simple custom Python function, all detector events that match these coordinates within a certain tolerance can be retrieved and visualized in an interactive plot. Traditionally, such an interactive plot represents only the XICs of the selected precursor and its fragments but this ignores the TIMS dimension. In contrast, with AlphaTims in this Jupyter notebook, we can easily provide heatmaps in both the LC and TIMS dimension for the precursor and all fragments, thereby illustrating the benefit of using TIMS data for peak capacity and interference removal. Using this extra information allows us to manually verify that the peptide of the spectral library is both quantitatively and qualitatively present in the diaPASEF data as well.

## Conclusion

The composition of a wide variety of (bio)chemical samples can be determined with LC-TIMS-QTOF, which acquires the intensity values of ions with billions of detector events that are convertible to chromatographic retention time, ion mobility, quadrupole *m*/*z* and TOF *m*/*z* values. While there are several tools that utilize this data for specialized applications, a generic software tool that is optimized for speed, usability and extensibility – thereby enabling community-driven developments – was lacking.

AlphaTims indexes unprocessed data in mere seconds, thereby making it equivalent to a sparse four-dimensional matrix. This allows to subsequently access the unprocessed data in milliseconds, regardless of the original complexity of the dataset. Due to this fast accession, AlphaTims also requires only milliseconds to provide interactive data visualizations along any dimension, including XICs, ion mobilograms, mass spectra, TICs or two-dimensional heatmaps. AlphaTims is easy to install and use on all major operating systems without requiring any computational expertise. It can be used as a stand-alone GUI, CLI or Python module and includes extensive help in the form of a README file, test data, a Python tutorial and a GUI manual. It is fully open-source with a minimal number of dependencies and is freely available under an Apache license at https://github.com/MannLabs/alphatims.

Due to the documented and freely available code base, AlphaTims can easily be modified or integrated in other projects. We ourselves for instance are already actively working on AlphaViz, a new software tool in the AlphaPept ‘ecosystem’ that visualizes identified peptides within raw data. Other examples of future modifications and applications include transferring parts of the indexing scheme to other vendors, a low-memory mode with optimized usage of HDF5 files, a multi-sample mode to directly compare different runs, or integration into automated quality control or feature finding pipelines.

## Acknowledgements

We would like to thank Sven Brehmer and Sascha Winter from Bruker Daltonics to explain the binary layout of analysis.tdf_bin files. Additional feedback from Nagarjuna Nagaraj and other Bruker Daltonics colleagues was also much appreciated. Finally, we are grateful for the feedback and support from within our own department, in particular Marvin Thielert, Andreas Brunner, Florian Meier, Igor Paron, Sophia Steigerwald and all members of the bioinformatics team and interest group.

## Abbreviations

CLI: (command-line interface)
DDA: (data-dependent acquisition)
DIA: (data-independent acquisition)
GUI: (graphical user interface)
JIT: (just-in-time)
LC: (liquid chromatography)
MGF: (Mascot generic file)
MS: (mass spectrometry)
MS/MS: (tandem mass spectrometry)
PASEF: (Parallel Accumulation–Serial Fragmentation)
Q: (quadrupole)
PyPi: (Python Package Index)
SPD: (samples per day)
TIC: (total ion current)
TIMS: (trapped ion mobility spectrometry)
TIMS: data format (tdf)
TOF: (time-of-flight)
XIC: (extracted ion chromatogram)

## Data availability

AlphaTims is fully open-source and is freely available with an Apache license at https://github.com/MannLabs/alphatims. The results in this manuscript were obtained with AlphaTims version 0.2.8. The mass spectrometry proteomics data have been deposited to the ProteomeXchange Consortium via the PRIDE^25^ partner repository with the dataset identifier PXD027359.

## Author contributions

SW implemented AlphaTims’ backend. EV implemented AlphaTims’ GUI. PS performed all sample preparation and acquisition. SW, EV, PS, MS and MM contributed ideas, performed testing and wrote the manuscript.

## References

1. Gabelica, V. et al. Recommendations for reporting ion mobility Mass Spectrometry measurements. Mass Spectrom. Rev. 38, 291–320 (2019).

2. Ridgeway, M. E., Lubeck, M., Jordens, J., Mann, M. & Park, M. A. Trapped ion mobility spectrometry: A short review. Int. J. Mass Spectrom. 425, 22–35 (2018).

3. Vasilopoulou, C. G. et al. Trapped ion mobility spectrometry and PASEF enable in-depth lipidomics from minimal sample amounts. Nat. Commun. 11, (2020).

4. Luo, M.-D., Zhou, Z.-W. & Zhu, Z.-J. The Application of Ion Mobility-Mass Spectrometry in Untargeted Metabolomics: from Separation to Identification. J. Anal. Test. 2020 43 4, 163–174 (2020).

5. Beck, S. et al. The impact II, a very high-resolution quadrupole time-of-flight instrument (QTOF) for deep shotgun proteomics. Mol. Cell. Proteomics 14, 2014–2029 (2015).

6. Fernandez-Lima, F., Kaplan, D. A., Suetering, J. & Park, M. A. Gas-phase separation using a trapped ion mobility spectrometer. Int. J. Ion Mobil. Spectrom. 14, 93–98 (2011).

7. Meier, F. et al. Online parallel accumulation–serial fragmentation (PASEF) with a novel trapped ion mobility mass spectrometer. Mol. Cell. Proteomics 17, 2534–2545 (2018).

8. Meier, F. et al. diaPASEF: parallel accumulation–serial fragmentation combined with data-independent acquisition. Nat. Methods 17, 1229–1236 (2020).

9. Łącki, M. K., Startek, M. P., Brehmer, S., Distler, U. & Tenzer, S. OpenTIMS, TimsPy, and TimsR: Open and Easy Access to timsTOF Raw Data. J. Proteome Res. 20, 2122–2129 (2021).

10. Yu, F. et al. Fast Quantitative Analysis of timsTOF PASEF Data with MSFragger and IonQuant. Mol. Cell. Proteomics 19, 1575–1585 (2020).

11. Lam, S. K., Pitrou, A. & Seibert, S. Numba: A LLVM-based Python JIT Compiler. in Proceedings of the Second Workshop on the LLVM Compiler Infrastructure in HPC - LLVM ‘15 (ACM Press).

12. Harris, C. R. et al. Array programming with NumPy. Nature vol. 585 357–362 (2020).

13. Folk, M., Heber, G., Koziol, Q., Pourmal, E. & Robinson, D. An Overview of the HDF5 Technology Suite and its Applications. in Proceedings of the EDBT/ICDT 2011 Workshop on Array Databases - AD ‘11 (ACM Press, 2011).

14. Wilhelm, M., Kirchner, M., Steen, J. A. J. & Steen, H. mz5: Space- and time-efficient storage of mass spectrometry data sets. Mol. Cell. Proteomics 11, O111.011379 (2012).

15. Askenazi, M., Ben Hamidane, H. & Graumann, J. The arc of Mass Spectrometry Exchange Formats is long, but it bends toward HDF5. Mass Spectrometry Reviews vol. 36 668–673 (2017).

16. Bhamber, R. S., Jankevics, A., Deutsch, E. W., Jones, A. R. & Dowsey, A. W. mzMLb: A Future-Proof Raw Mass Spectrometry Data Format Based on Standards-Compliant mzML and Optimized for Speed and Storage Requirements. J. Proteome Res. 20, 172–183 (2020).

17. Cottam, J. A., Lumsdaine, A. & Wang, P. Abstract rendering: out-of-core rendering for information visualization. in Visualization and Data Analysis 2014 (eds. Wong, P. C., Kao, D. L., Hao, M. C. & Chen, C.) vol. 9017 90170K (SPIE, 2013).

18. Strauss, M. T. et al. AlphaPept, a modern and open framework for MS-based proteomics. bioRxiv 2021.07.23.453379 (2021) doi:10.1101/2021.07.23.453379.

19. Kulak, N. A., Pichler, G., Paron, I., Nagaraj, N. & Mann, M. Minimal, encapsulated proteomic-sample processing applied to copy-number estimation in eukaryotic cells. Nat. Methods 11, 319–324 (2014).

20. Bache, N. et al. A novel LC system embeds analytes in pre-formed gradients for rapid, ultra-robust proteomics. Mol. Cell. Proteomics 17, 2284–2296 (2018).

21. Eisenstat, S. C., Gursky, M. C., Schultz, M. H. & Sherman, A. H. Yale sparse matrix package I: The symmetric codes. Int. J. Numer. Methods Eng. 18, 1145–1151 (1982).

22. Mckinney, W. pandas: a Foundational Python Library for Data Analysis and Statistics. http://pandas.sf.net.

23. Egertson, J. D., MacLean, B., Johnson, R., Xuan, Y. & MacCoss, M. J. Multiplexed peptide analysis using data-independent acquisition and Skyline. Nat. Protoc. 10, 887–903 (2015).

24. Muntel, J. et al. Surpassing 10 000 identified and quantified proteins in a single run by optimizing current LC-MS instrumentation and data analysis strategy. Mol. Omi. 15, 348–360 (2019).

25. Perez-Riverol, Y. et al. The PRIDE database and related tools and resources in 2019: improving support for quantification data. Nucleic Acids Res. 47, D442–D450 (2019).

